# Extracting active modules from multilayer PPI network: a continuous optimization approach

**DOI:** 10.1101/211433

**Authors:** Dong Li, Zexuan Zhu, Zhisong Pan, Guyu Hu, Shan He

## Abstract

Active modules identification has received much attention due to its ability to reveal regulatory and signaling mechanisms of a given cellular response. Most existing algorithms identify active modules by extracting connected nodes with high activity scores from a graph. These algorithms do not consider other topological properties such as community structure, which may correspond to functional units. In this paper, we propose an active module identification algorithm based on a novel objective function, which considers both and network topology and nodes activity. This objective is formulated as a constrained quadratic programming problem, which is convex and can be solved by iterative methods. Furthermore, the framework is extended to the multilayer dynamic PPI networks. Empirical results on the single layer and multilayer PPI networks show the effectiveness of proposed algorithms.

Availability: The package and code for reproducing all results and figures are available at https://github.com/fairmiracle/ModuleExtraction.

## 1 Introduction

Identifying functional modules from protein-protein interaction (PPI) networks is one of the central challenge in network biology [1]. The modules (or subnetworks) represent certain interacted components of the whole network to exert some functions, which may shed light to the complex mechanisms behind living organisms. Modular structure in various PPI networks has been reported in [2, 3, 4, 5], and community detection techniques [6, 7] have been applied in a wide range of biological networks, including PPI networks.

Though the topology of a biological network does not always precisely reflects the function or even disease-determined regions [8], which are the real concerns in biology. The topological modules and functional modules may have some overlapped components, but still vary from definition. To better address these concerns, the **active module** identification, which aims to find connected regions of the network showing striking changes in molecular activity or phenotypic signatures that are associated with a given cellular response in biological networks [9], has become an important issue in network biology.

Ideker et al. [10] proposed a general method to discover active connected regions by given network with scores on each node, which reflects the changes over particular conditions. The seminal work [10] not only defined an optimization problem on a weighted molecular interaction networks, which incorporates basic network structure with high-throughput omics data, but also proposed simulated annealing algorithm to solve it. Some following researches improved this basic active module identification from many aspects, resulting into a series of heuristic based methods [11, 12, 13, 14] and integer programming based methods [15, 16]. A comprehensive review [9] listed several advances in this area.

However, in the existing works of active modules identification, node [15] or edge weights [13] (or both [17]) weight too much in original active module identification objective, which leads to modules without topological properties other than connectivity. In other words, current active modules identification algorithms tend to categorize several unrelated nodes in terms of their interactions pattern.

But some earlier results derived from pure topological structure, such as communities [6] or motifs [18], also make sense of biological meaning [19]. The communities are important since they may correspond to functional units [20]. As a recent example of 2016 Disease Module Identification DREAM Challenge [21], the participators were asked only to use network structural information to find disease modules, which were evaluated by pre-defined genome-wide association study (GWAS) sets. Quite a few teams only adopted basic communities detection methods which maximize the modularity [22, 23] also achieved good performance. As a community effort, basic modules detection algorithms which only consider the topology of biological networks are proven to be effective as well.

In this work, we try to combine these two parallel lines of research, active modules identification and communities detection, to extract modules from PPI networks. Specifically, we consider the objective of representing an active module is composed of two parts: the topological part and the active part. By combining these two parts, we can reveal functional change and regulatory or signaling mechanisms. The topological property of a module is derived from graph partitioning, and the active property is highlighted by the higher expected average node score. As a result, we formulate the active modules identification on connected PPI network as a constrained quadratic programming problem, which can be solved efficiently by iterative methods. We also show that this method can be extended to multilayer networks, by conducting the same algorithm on an aggregation of multiple networks. The package and code for reproducing all results and figures are available at https://github.com/fairmiracle/ModuleExtraction.

## 2 Method

### 2.1 Optimization on single-layer PPI network

We aim to extract a module from a weighted network with both topological and active features. The problem is formally defined as:

#### Problem 1.

*Given a weighted graph G* = (*V*, *E*, *W*) *where w*_*ij*_ *is the weight between vertex i and j, vertices weights z*_*i*_ *for each i* ∈ *V, find a connected subnetwork S* = (*V*_*S*_, *E*_*S*_) *of G with significant separation from the rest in both edges and vertices weight.*

The separation from topological perspective has been intensively studied in graph theory. We thus start with the logic of spectral clustering (partitioning) on the graph (network) *G.* The *cut* for two disjoint subsets *A* and its complement 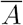 on *G,* defined as

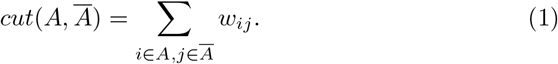

The basic criteria to measure the quality of a partition on graph *G* is to minimize 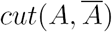 over all possibilities of *A.* But there exists a trivial case when *A = V* or *A* = Ø. Even when there is only one node or very few nodes in *A,* the 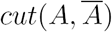 is small, which is not preferred. Several modifications based on *cut* were proposed, such as the *ratio cut* [24]:

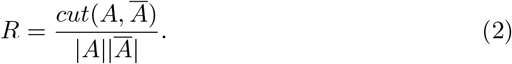

Zhao et al. [25] proposed the community extraction framework based on *ratio cut,* incorporating with the assumption that the interactions in the extracted module should be denser. The same intuition had been formulated as the well-known *modularity* [22], which also measures the quality of a partition instead of certain extraction module. Adopting the actual-minus-expected paradigm on the average node score, we define the following objective to be minimized:

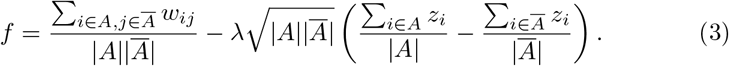

The first part is the same as *ratio cut,* which is supposed to be minimized from the topological perspective. And 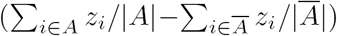 in the second term is the gap between the expected node score in the module and in the rest, which is supposed to be maximized (equivalent to minimize the negative) from the active consideration. We also add the penalty 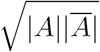 to this part, when 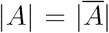 the penalty has a maximal value, which is consistent with the denominator of *ratio cut.* Parameter λ is the used to balance the consideration between edge part and node part. Without prior knowledge or preference on the two parts, we suggest the default value λ = 1.

Being similar to Luxburg [26] (section 5, Graph cut point of view), we define the vector x = (*x*_1_,*x*_2_,…, *x*_*n*_)^⊤^ with entries:

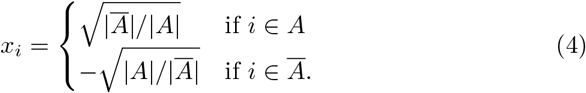

Note the following two facts which are the same as in [26]:

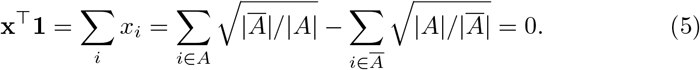

and

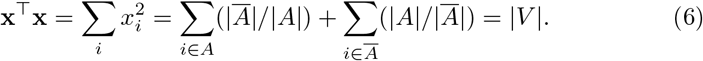

We also have

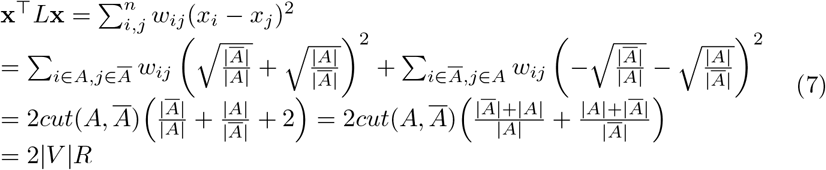

and

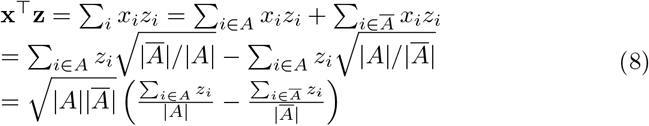

Combining equation (7), (8), and conditions (5) and (6), we can rewrite the objective (3) into the matrix form:

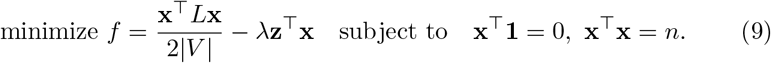

Since the objective function is smooth and differentiable, we use a projected gradient [27] approach to find the local optimum, which works as follow. The gradient is used to determine the primary direction, followed by orthogonalization to satisfy the condition (5) and normalization step to satisfy the condition (6). Following [28] we claim that any limit point of sequence {x^(*k*)^} generated by this procedure is a stationary point of (3). When *L* is positive semi-definite, the problem is convex thus a local optimum is also the the global optimum.

After getting the target set *A*, it may comes to case that the candidates nodes are not connected, due to the relaxation on x, which was supposed to be 0-1 vector. We conduct a connected components finding (CCF) operation on *A* as in [29] to extract large component. The existence of giant component in a random set of vertices from a graph is supported by the Erdös–Rényi model [30]. In summary, the procedure is formally described as Algorithm 1.

**Algorithm 1:**
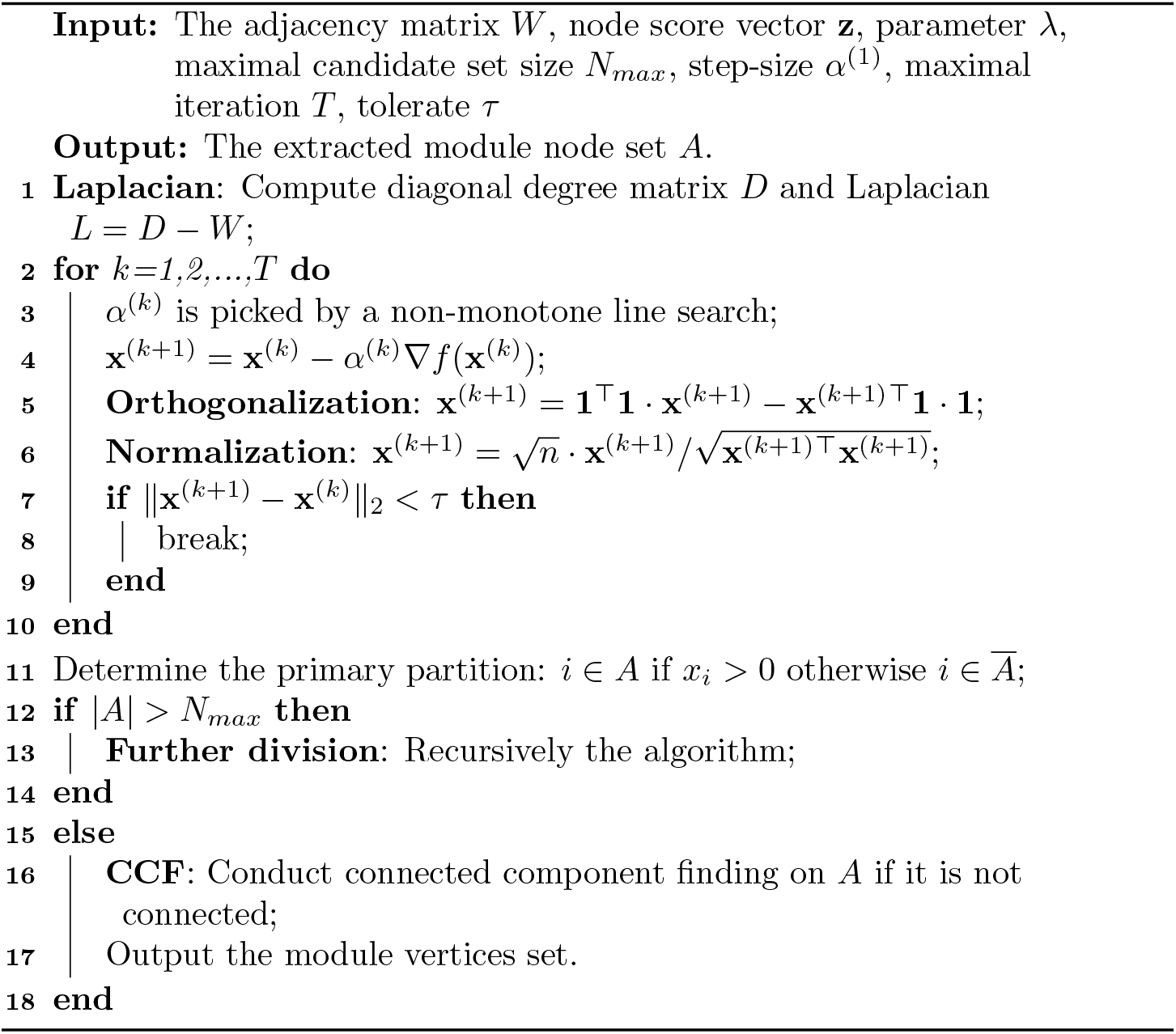
Module extraction via continuous optimization.

For a large network, we may need to execute Algorithm 1 multiple times, to get proper number of modules. There are two strategies: recursively extraction and components finding. The former works for dense interacted networks, which mines module one by one and remove the identified module from background network. And the latter aims to leverage the relatively small modules from the candidate set, which avoids wasting isolated nodes when the interactions are not enough.

### 2.2 Aggregation on multilayer PPI network

The community structure has been explored on multilayer connected graphs. Various algorithms have been proposed to detect the communities on multilayer graphs, see the review [31]. One of the nature idea is to extend the concept of modularity to multilayer case, as [32] did. Taking the layer into consideration, the matrix calculation in original modularity was transformed into tensor computation. The underlying assumption of this extension is that each layer shares similar structure, and the word “multiscale” in the title of [32] also indicates that each network shares similar structure.

However, in some real applications, the multiple layers can be diverse. The sub-challenge 2 of Disease Module Identification DREAM Challenge [21] required find modules across multiple networks, including two protein-protein interaction networks, one signaling network, one co-expression network, one cancer network and one homology network. In such a multilayer network system, the size and topology are diverse with all nodes aligned. Due to the fact that there be very less shared edges among different networks, even no single edge shared by all networks, we can highlight the majority of presence of edges by adopting the idea of aggregated graph or consensus graph. An aggregated graph is simply defined by the sum of all edge weight matrices, followed by proper cut-off to remove low valued entries. The intuition behind this aggregating is to enhance the concurrent edges across more layers and remove layer-specific edges. As a result, the aggregated graph is supposed to encode the conservation information of multiple layers. And the module (defined by the solution of (3)) extracted from this graph represents conserved subnetwork with maximal node activities across multiple layers. When each layer is a snapshot of a dynamic system, the extracted modules can be viewed as time-invariant subnetworks, which may be related to persistent biological processes.

The complete procedure to mine modules from multiple networks *G*_1_, *G*_2_, …, *G*_*n*_ is describe as Algorithm 2.

**Algorithm 2:**
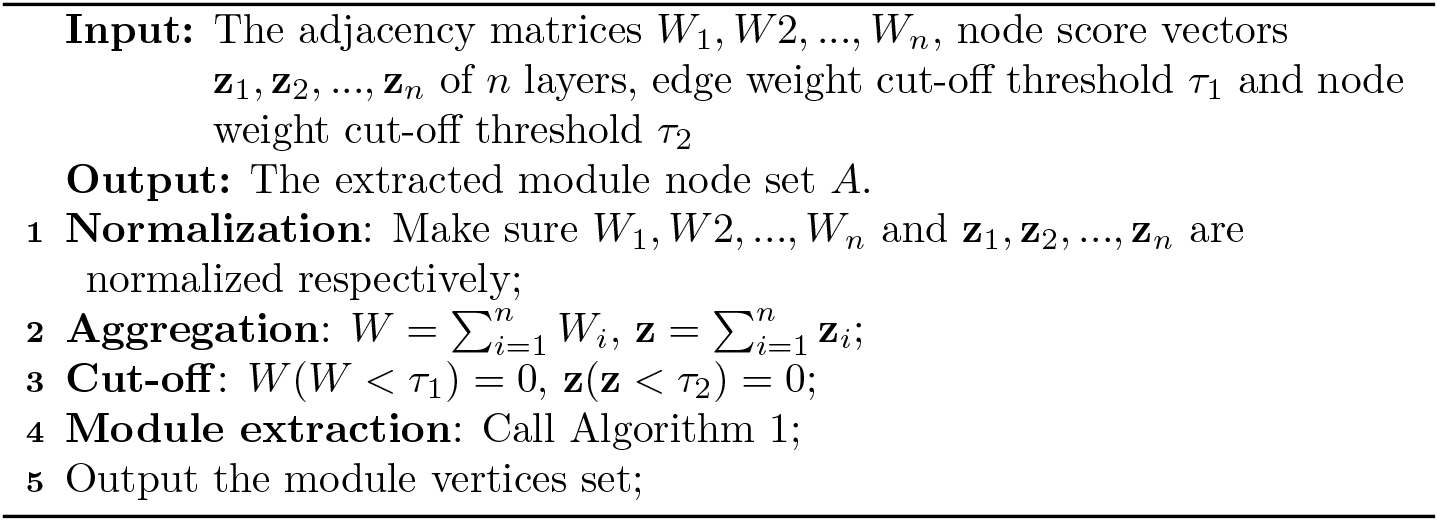
Module extraction via continuous optimization on multilayer network.

## 3 Results and Discussion

### 3.1 Toy example

We first compare the proposed method with *ratio cut* minimization, which actually solves the graph partitioning without using the node score:

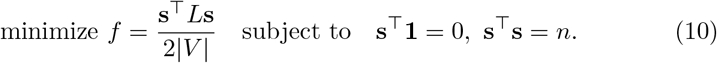

According to Rayleigh-Ritz theorem [26, 33], the solution is directly given by the eigenvector corresponding to the second smallest eigenvalue of *L*, denoted as u. The nodes of extracted module are selected according to the sign of entries in u, i.e. *i* ∈ *A* if *u*_*i*_ > 0. We denote this method as spectral algorithm in the following discussion.

We compare the spectral algorithm with Algorithm 1 on the widely used Zachary karate club network, as Figure 1 (a) shows. Red nodes mean the members in the target module. The simulated node score of all nodes are randomly generated via **z** ~ [0, 1], but with higher expectation in target module members as **z** ~ [0.5, 1]. We can see that spectral algorithm (see Figure 1c) not only failed to find the important node ‘34’ in the target module, but also wrongly include two nodes ‘1’ and ‘17’ from the opposite module. Furthermore, node ‘1’ is considered to play a core role in the opposite module. The proposed method (see Figure 1c) successfully found all the nodes but introduced additional node ‘3’, which also interacts with other nodes in the target module. In Figure 1c the node size indicates the assigned score, and the result validates the purposed of (9), which combines modular structure and high scored nodes. In other words, not all high scored nodes are included, such as the connected and high-scored nodes ‘20’, ‘1’, ‘13’ and ‘22’.

**Figure 1:**
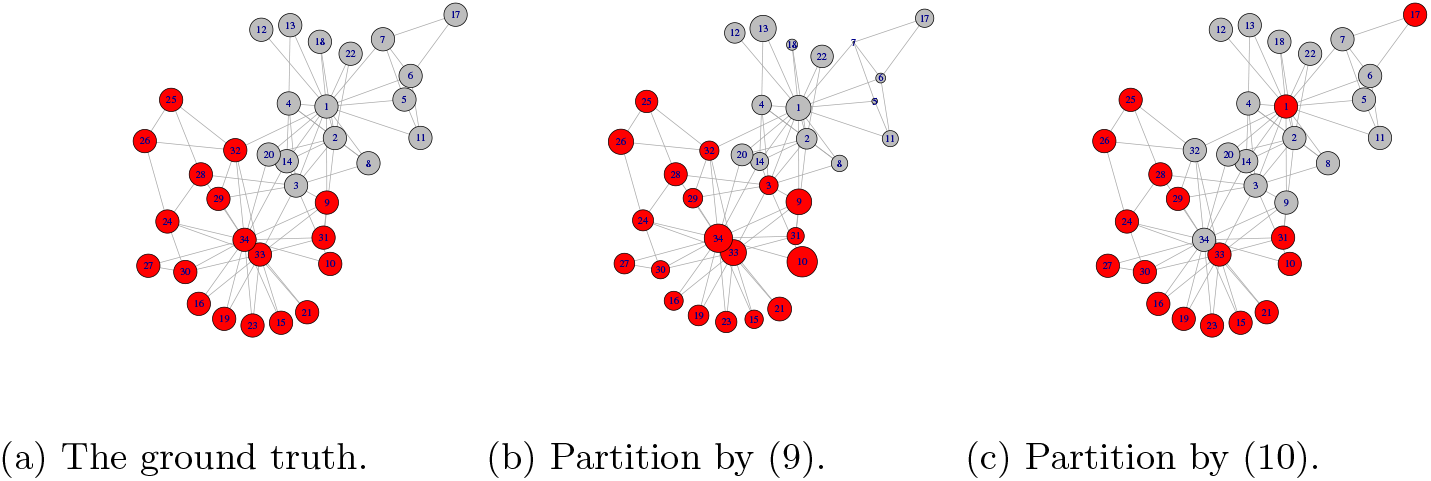
Comparison of the ground truth with partition by proposed method (9) and spectral algorithm (10), on the widely used Zachary karate club network. Red nodes mean the members in the target module. And node size indicates the simulated node score.

### 3.2 Single layer PPI network

In order to check the biological relevance of identified modules by the proposed algorithm, we apply it on the real world protein-protein interactions (PPI) network. The human PPI network and p-values derived from differential expression and survival analysis [34] come from package BioNet [35], which implements the integer programming based active module identification algorithm [15] as well as a heuristics algorithm. The heuristics algorithm can be viewed as an alternative of the Cytoscape plugin jActiveModules [10]. Being different from the objective (3), both the exact approach [15] and the heuristics method [10] aim to collect as more high-scored and connected nodes but the exact approach relies on external library CPLEX.

We compare the heuristics method from BioNet and Algorithm 1 (using default parameters: λ = 1, *α* = 0.01, τ = 1*E* – 9, *T =* 100, *N*_*max*_ = 100) on the same PPI network with 2559 nodes and 7788 edges, and the nodes are scored by the Beta-Uniform-Mixture (BUM) model [15] based on expression p-values. BioNet identifies a module with 37 nodes and 44 edges, and the Algorithm 1 identifies a module with 54 nodes and 54 edges. And there are 20 nodes in common, most of which are high-scored nodes. In order to show the difference directly, we plot both modules in one figure, with different colors, as Figure 2 shows. We can see that in the both modules, there are some low-scored nodes such as SMAD2, LYN, CDC2, NME1, serving as bridge nodes which connect server component. But the module identified by Algorithm 1 includes more low-scored nodes and their interactions.

The identified modules are enriched by biological progresses and pathways related to the experimental settings. Since these two modules have different size, we use the false discovery rate (FDR) of enriched KEGG pathways to show the significance of enrichment analysis. Table shows the top 5 pathways enriched by both modules, and we can see find more significant results. Furthermore, all pathways are related to cancer, which is consistent with the data source [34].

**Figure 2:**
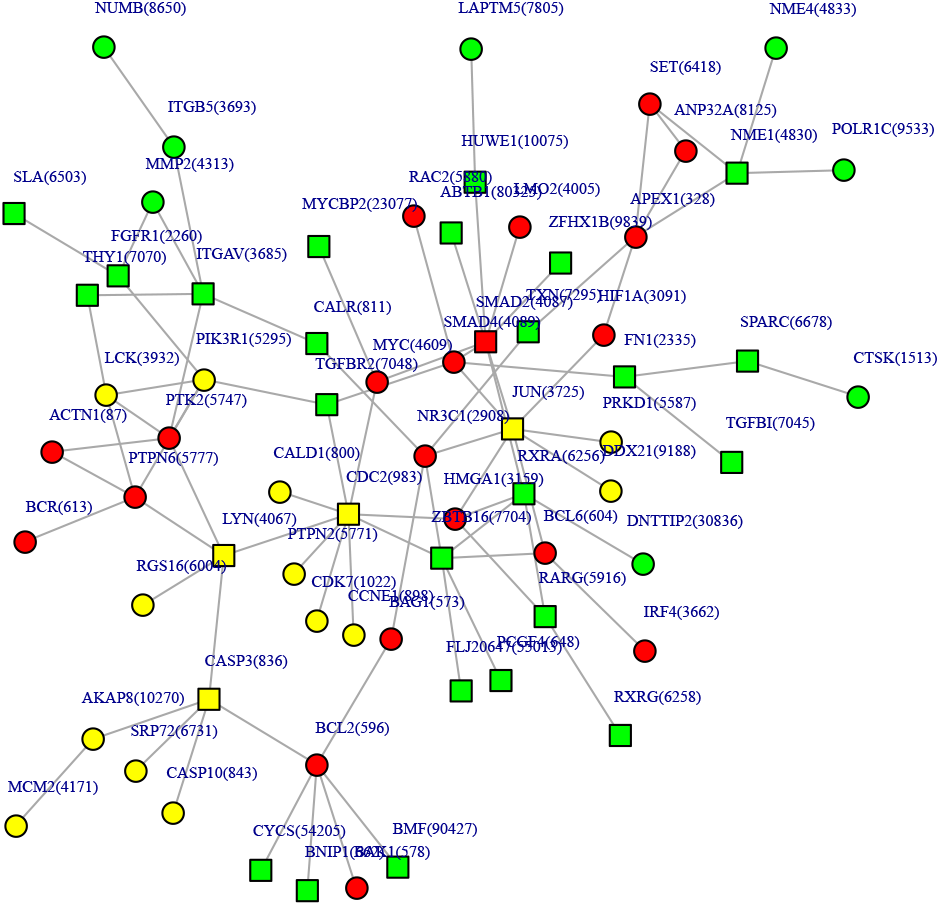
The identified modules by BioNet and Algorithm 1. The red nodes are shared by both methods, yellow for BioNet and green for Algorithm 1. The shape of nodes indicates score, squares indicate negative scores and circles for positive.

### 3.3 Multilayer dynamic PPI networks

Multilayer dynamic PPI networks (DPPI or DPIN) are normally constructed by integrating static PPI network and a time-course gene expression data [36, 37, 38]. Each layer in a DPPI network represent a specific PPI of a time point, and the whole DPPI is supposed to model the dynamic properties of protein interactions. While the active modules across all layers can be viewed as complementary to those identified from static PPI [39], due to the time course data integration. Here we adopt the DPPI construction method by [38, 36], where the topological structure of each layer is determined by both the static yeast PPI network and the gene expression data at that time point. In addition, the node activity of each layer simply measured by the expression values. The processed gene expression data as well as DPPI construction can be found in [36].

First of all, we try to demonstrate the advantage of multilayer aggregation by comparing the result from the consensus graph and each single layer. We evaluate the result by f-measure [40], which assesses the similarity between identified modules (protein complexes) *C*_*i*_ ∈ *P* and reference complexes *C*_*j*_ ∈ *B.* We claim *C*_*i*_ and *C*_*j*_ match if *v*_*ij*_ ≥ 0.25:

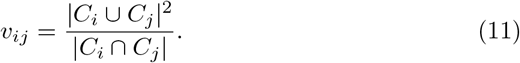

F-measure is defined based on precision and recall:

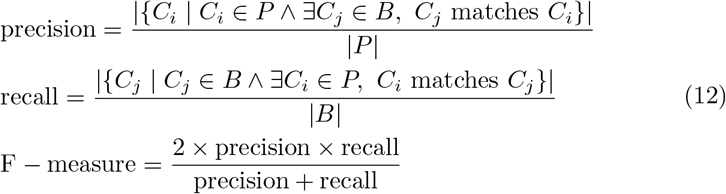

Two widely used reference complexes CYC2008 [41] and MIPS [42] are chosen as gold standard sets *B.* Furthermore, we use SPICi [43] as baseline algorithm which was designed for detecting non-overlapping complexes from static PPI. Also note algorithms for detecting overlapping complexes such as ClusterONE [44] and TS-OCD [36] would achieve higher F-scores since part of proteins might appear frequently in different modules, which increases the chance to be captured by reference sets according to (12).

Figure 3 shows the overall result of modules from each layer, consensus graph and the baseline algorithm SPICi [43]. The purpose of this figure is twofold: identifying modules on the consensus graph is superior to each single one with the same algorithm and Algorithm 2 could achieve comparable result with other methods in terms of accuracy. Specifically, Algorithm 2 tends to achieve higher precision but lower recall compared with SPICi [43]. Possible reason for failing to recall as more complexes lies at the CCF operation in Algorithm 1, which may miss some proteins in the reference set.

**Figure 3:**
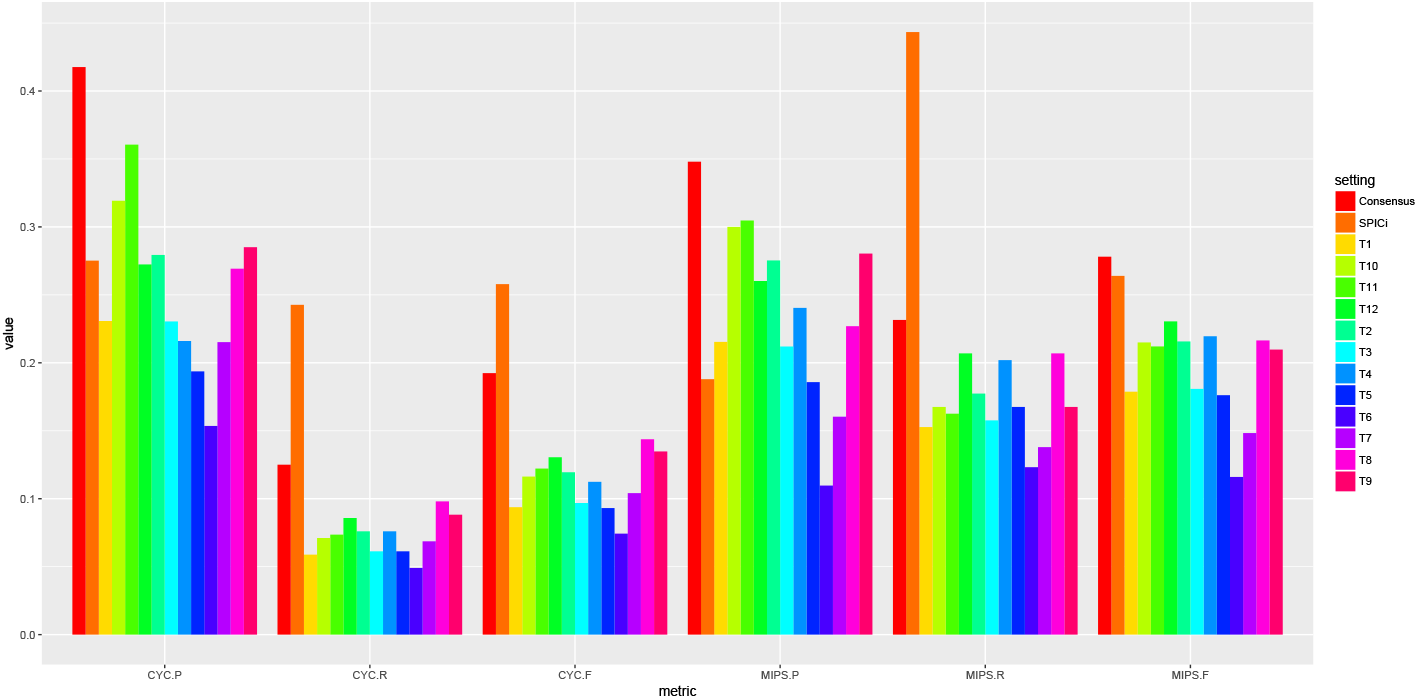
The F-measure of identified complexes from each time point layer, consensus graph and the baseline algorithm. ‘CYC-P’ means the precision evaluated by CYC2008 [41] and ‘MIPS-R’ means the recall evaluated by MIPS [42]. ‘F’ is the F-measure defined as in (12).

In order to validate the biological meaning of the identified complexes, we conduct enrichment analysis with STRING [45], which provides interfaces to access the annotation resources as well as enrichment result. Furthermore, we compare the results of Algorithm 1 and SPICi [43] in a statistical way, and count the number of complexes which are significantly enriched by at least one GO term (biological processes and KEGG pathways) at a given FDR (≤0.05) cutoff. From table 2 we can see that Algorithm 2 found relatively larger complexes on average, and the resulted complexes are enriched by more KEGG pathways. The ratio of meaningful complexes is also higher than the result from static PPI network. See the complete enrichment analysis of these modules from two algorithms in supplementary table SS1 and SS2.

**Table 1:** The top 5 enriched KEGG pathways of the modules identified by Algorithm 1, and heuristics algorithm which targets at a group of high-scored and connected nodes, on single human PPI network.

| Algorithm 1 |  |  | Heuristics algorithm |  |  |
| --- | --- | --- | --- | --- | --- |
| ID | Description | FDR | ID | Description | FDR |
| 5200 | Pathways in cancer | 5.05E-16 | 5200 | Pathways in cancer | 1.23E-9 |
| 5222 | Small cell lung cancer | 5.36E-9 | 5210 | Colorectal cancer | 1.23E-9 |
| 5210 | Colorectal cancer | 1.03E-8 | 5161 | Hepatitis B | 4.77E-7 |
| 5205 | Proteoglycans in cancer | 1.18E-8 | 4110 | Cell cycle | 5.43E-6 |
| 5202 | Transcriptional misregulation in cancer | 1.72E-8 | 4520 | Adherens junction | 9.32E-6 |

**Table 2:** Overview of the resulted complexes of Algorithm 2 and SPICi [43]. *#Items* means the total GO terms of all complexes and *Ration* is defined as the proportion of complexes which are significantly enriched by at least one GO term at a given FDR (≤0.05).

| Method | Modules | Avg. size | Enrichment |  |  |
| --- | --- | --- | --- | --- | --- |
|  |  |  |  | BP | KEGG |
| SPICi | 298 | 4.21 | #Items | 8482 | 599 |
|  |  |  | Ratio | 0.81 | 0.72 |
| Algorithm 2 | 277 | 5.06 | #Items | 7114 | **1088** |
|  |  |  | Ratio | 0.81 | 0.88 |

## 4 Conclusion

This paper presents a continuous optimization approach to extract active modules from PPI networks. The method tries to optimize the objective combining both topological and active properties of a module, which reflects the functional organization mechanism in a complex system. We also generalize the proposed method to the multilayer dynamic PPI networks, using an aggregation of multiple single layers. Empirical studies shows the advantages of the optimization approach as well as the aggregation on multilayer networks.

